# Landscape of human miRNA variation and conservation using <u>A</u>nnotative <u>D</u>atabase of <u>miR</u>NA <u>E</u>lements, ADmiRE

**DOI:** 10.1101/177170

**Authors:** Ninad Oak, Rajarshi Ghosh, Sharon E. Plon

## Abstract

MicroRNAs (miRNAs) are the most abundant class of non-coding RNAs that regulate expression of >60% genes and are frequently deregulated in many human diseases. Sequence variants in miRNAs are expected to have a high impact on miRNA function. However, the lack of miRNA variant annotation and prioritization guidelines has hampered this analysis from whole genome/exome sequencing (WGS/WES) studies. Through the development of an <u>A</u>nnotative <u>D</u>atabase of <u>miR</u>NA <u>E</u>lements, ADmiRE workflow, we re-annotated the publicly available population dataset of gnomAD 15,596 WGS and 123,136 WES and describe 26,094 precursor-miRNA variants. AdmiRE annotates twice the miRNA variants predicted by existing tools which prioritize variation relative to protein coding regions. We provide the allele frequency distribution of miRNA variation which is comparable to variation in exonic regions. This distribution is similar for miRNAs located in the intragenic and intergenic genomic context. Moreover, ‘high confidence’ miRNAs (designated by miRBase) harbor less variation (the majority contributed by rare variants) compared with the remaining miRNAs. We identify 279 miRNAs highly constrained with little or no variation in gnomAD. We further describe the evolutionary conservation of miRNAs across 100 vertebrates and identify 434 highly conserved miRNAs. We demonstrate that these constraint and conservation metrics (now incorporated into the ADmiRE workflow) characterize miRNAs previously implicated in human diseases. In conclusion, through the development of ADmiRE, we comprehensively analyze the landscape of miRNA sequence variation in large human population datasets and provide miRNA vertebrate conservation scores to aid future studies of miRNA variation in human diseases.

## Introduction

MicroRNAs (miRNAs) are small non-coding RNAs, 18-25 nucleotides in length, that regulate the expression of greater than 60% of genes by complementarily binding target mRNAs (Friedman et al. 2009; Bartel 2009). Over the last decade, miRNAs have been extensively profiled for their expression across human diseases leading to the development of several miRNA-targeted therapeutics (Rupaimoole and Slack 2017; Almeida et al. 2011). A number of underlying mechanisms for aberrant miRNA expression have been proposed including genomic amplifications or deletions, single nucleotide variation and transcriptional regulation (Chan et al. 2011). For example, germline or somatic variants in miRNA transcripts are associated with several cancers such as breast cancer (Shen et al. 2009, 2010; Li et al. 2009), leukemia (Calin and Croce 2006; Kotani et al. 2010; Calin et al. 2005) and pancreatic cancer (Zhu et al. 2009). Sequence variation in miRNA intermediate transcripts, such as in primary-and precursor-miRNA sequences, can interfere in the miRNA biogenesis pathway (Zeng and Cullen 2003). However, there have been few systematic reports describing and analyzing genetic sequence variation across miRNAs in large population datasets which is imperative to fully understand the role of miRNA variation in human diseases.

Existing studies for miRNA sequence variation have been limited by small sample size and small subsets of the human miRNAs assessed. An early study identified a total of 527 precursor miRNA variants by analyzing 720 miRNAs from the 1000 Genomes Project, Phase I (Carbonell et al. 2012) dataset. Other databases such as miRNASNP and polymiRTS report ∼500 single nucleotide polymorphisms (SNPs) in miRNA regions using the dbSNP137 database (Gong et al. 2012; Bhattacharya et al. 2014). However, a growth in miRNA sequence identification (miRBase v21: 1,881 precursor, 2,815 mature miRNAs) and in large whole genome sequencing (WGS) datasets warrants comprehensive analysis of miRNA variation (Kozomara and Griffiths-Jones 2011).

Interpretation of noncoding sequence variants from large whole genome sequencing studies has proved to be challenging due to lack of well-defined functional domains, precise biological consequence predictions and poor understanding of sequence constraint in large human reference datasets (Ritchie et al. 2014). miRNAs as an important class of regulatory noncoding RNAs are not well annotated by many commonly used variant annotation tools such as variant effect predictor (VEP) (McLaren et al. 2016), ANNOVAR (Wang et al. 2010) or SnpEff (Cingolani et al. 2012) and/or the programs’ associated plugins provide limited information regarding miRNA variants (McLaren et al. 2016). Many of these tools often prioritize variants relative to protein-coding genes over noncoding RNA variants, such as those in miRNAs. Furthermore, interpretation of protein-coding variation often relies on predictors of deleteriousness of nonsynonymous variants such as PolyPhen (Adzhubei et al. 2010) and SIFT (Kumar et al. 2009) based on evolutionary conservation or on sequence constraint metrics developed using large human population datasets such as ExAC (Lek et al. 2016; Samocha et al. 2014, 2017). In comparison, establishing similar sequence constraint and conservation metrics for well-defined noncoding RNAs such as miRNAs is expected to improve variant interpretation in these regions. However, the only dedicated miRNA variant annotation tool, miRVaS (Cammaerts et al. 2015), does not include the sequence constraint metrics nor the biological annotations such as expression, downstream targets, and disease associations.

In this study, we describe a novel miRNA variation module entitled <u>A</u>nnotative <u>D</u>atabase for <u>miR</u>NA <u>E</u>lements, ADmiRE (https://github.com/nroak/ADmiRE) for variant calling and annotation of variants in human miRNAs that can be easily integrated into established annotation workflows of variant files (VCF, TSV, etc.) or used independently from BAM files. ADmiRE annotates twice the variants as within miRNAs compared to VEP. Using this new workflow, we analyze existing publicly available datasets including gnomAD containing 123,136 WES and 15,496 WGS samples which allow us to explore the domain level and miRNA level sequence constraints for miRNA variation, similar to previous work done on protein coding genes (Lek et al. 2016). We also perform parallel analysis of miRNA sequence conservation across 100 vertebrates and incorporate these analyses into ADmiRE. In sum, development of a dedicated efficient miRNA variant annotation tool which includes sequence constraint and conservation information for human miRNAs in combination with existing miRBase annotations can guide the prioritization and identification of biologically relevant miRNA variation in control and disease datasets.

## Results

### Development of Annotative Database of miRNA Elements (ADmiRE)

We developed an <u>A</u>nnotative <u>D</u>atabase of <u>miR</u>NA <u>E</u>lements or ADmiRE, that provides a rich resource for the interpretation of human miRNA variation (see Methods). ADmiRE contains information about the location of the variant in miRNA sequence domains (Figure 1A), base-pair conservation scores and based on the work described in this paper, also includes whether a miRNA is constrained in gnomAD and/or conserved across 100 vertebrates.

**Figure 1.**
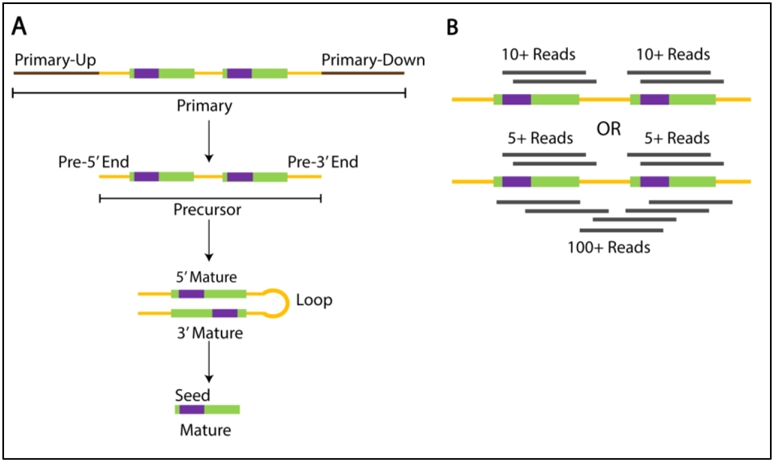
Schematic of miRNA sequence domains and “high confidence” annotations. **A.** miRNA Sequence domains as annotated in ADmiRE pipeline. Precursor (yellow) and mature (green) domains as defined by miRBase, Primary (brown) domains defined as 100bp flanking regions, and Seed (purple) domains defined as 2-8bp of mature domains. **B.** Definition of ‘high confidence’ miRNAs based on the evidence from miRNAseq experiments as described by miRBase v21.

ADmiRE further integrates the biologically relevant information regarding ‘high confidence’ annotations (based on miRNA expression studies) from miRBase (Figure 1B) (Kozomara and Griffiths-Jones 2014), experimentally validated targets of miRNAs and their human disease associations, making it a powerful tool for the interpretation of variant impact in the most updated list of miRNAs from miRBase. We integrated ADmiRE annotations into the widely used single nucleotide variant calling workflow from Genome Analysis Toolkit (GATK) to call miRNA variation from WGS/WES alignments (BAM files) (McKenna et al. 2010; Van der Auwera et al. 2013) (Figure 2A, 2B). ADmiRE workflow can be accessed freely to recall miRNA variants from BAM files or to annotate an existing set of variants (VCF, MAF, and other formats) with miRNA specific annotations using a standalone program or integration into VEP pipeline (see Data Access).

**Figure 2.**
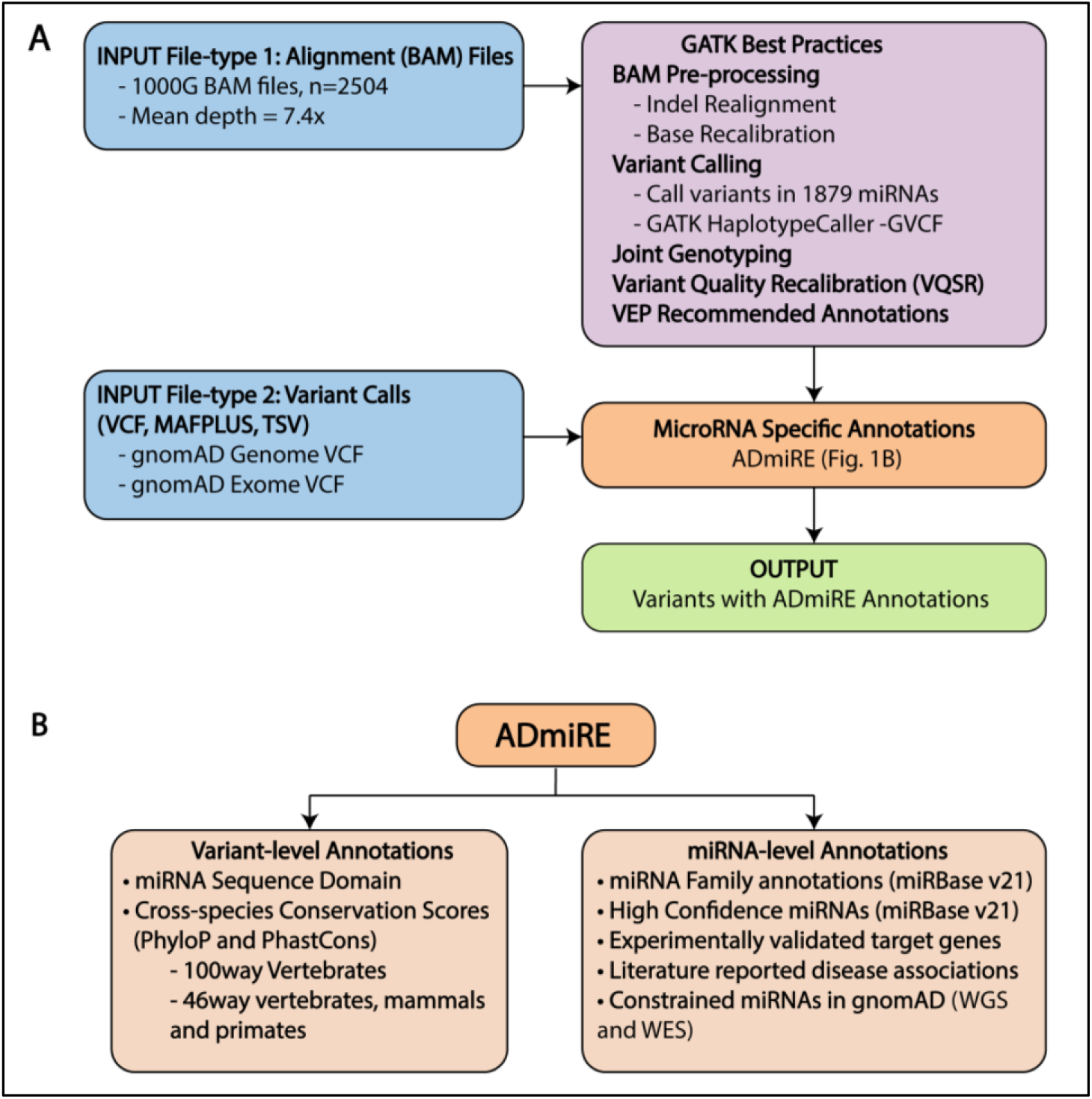
Annotative Database of miRNA Elements, ADmiRE, Workflow. **A.** Variant calling pipeline based on GATK Best Practices workflow that takes a sequence alignment (BAM) file as an input, preprocesses it and performs joint variant calling using GATK HaplotypeCaller. A second input option with pre-called variants is available. Variants are annotated using ADmiRE database and an annotated output file is generated. **B.** Description of ADmiRE annotations that are added for every variant in a miRNA region.

Initial application of this workflow was demonstrated using the publicly available high-quality sequence alignments and VCFs released under the 1000 Genomes Project (1000G) including analysis of a unique subset of 20 individuals from 1000G project where each sample was sequenced by three of the most commonly used sequencing platforms, namely, high coverage WES (mean depth = 65.7x), high coverage WGS (mean depth >30x) and low coverage WGS (mean depth =7.4x) (see Methods). Not surprisingly, the high coverage WGS platform captured >99% of miRNAs in miRBase v21 at a high read depth (>30X) while WES captured about ∼54% miRNAs with read depth greater than 15X (Supp. Fig. 1). We provide ADmiRE analysis of human miRNA variation in the publicly available population datasets 1000G and UK10K which are available in Supplementary Data. In the following sections of this paper, we describe miRNA variation analysis of the large ExAC and gnomAD datasets.

### ADmiRE uncovers additional miRNA variation in existing datasets

We compared miRNA annotations from commonly used variant annotation tools with ADmiRE. Most existing tools use a set of pre-defined criteria for consequence prioritization which is currently optimized for protein-coding regions. Annotation of variants in ExAC dataset (60,706 WES samples) using ADmiRE workflow identified a total of 6,639 mature domain variants 2,764 of the 2,964 mature domain variants previously annotated by Variant Effect Predictor (VEP) (Figure 3A). Of the 200 variants missed by ADmiRE, only 27 variants mapped to miRNA regions when verified using UCSC genome browser. Thus, ADmiRE identified approximately twice the number of mature miRNA variants compared to VEP while missing only 0.4% of verified miRNA variants. We found that the additional 3,875 variants detected by ADmiRE were most frequently annotated by VEP as ‘non_coding_transcript_exon_variant’ (50%), the category describing all non-coding RNAs including precursor miRNA domains, and ‘intron_variant’ (29%) (Figure 3B). Overall, VEP prioritizes the annotation for intronic protein-coding domains over mature miRNA and other non-coding RNA domains.

**Figure 3.**
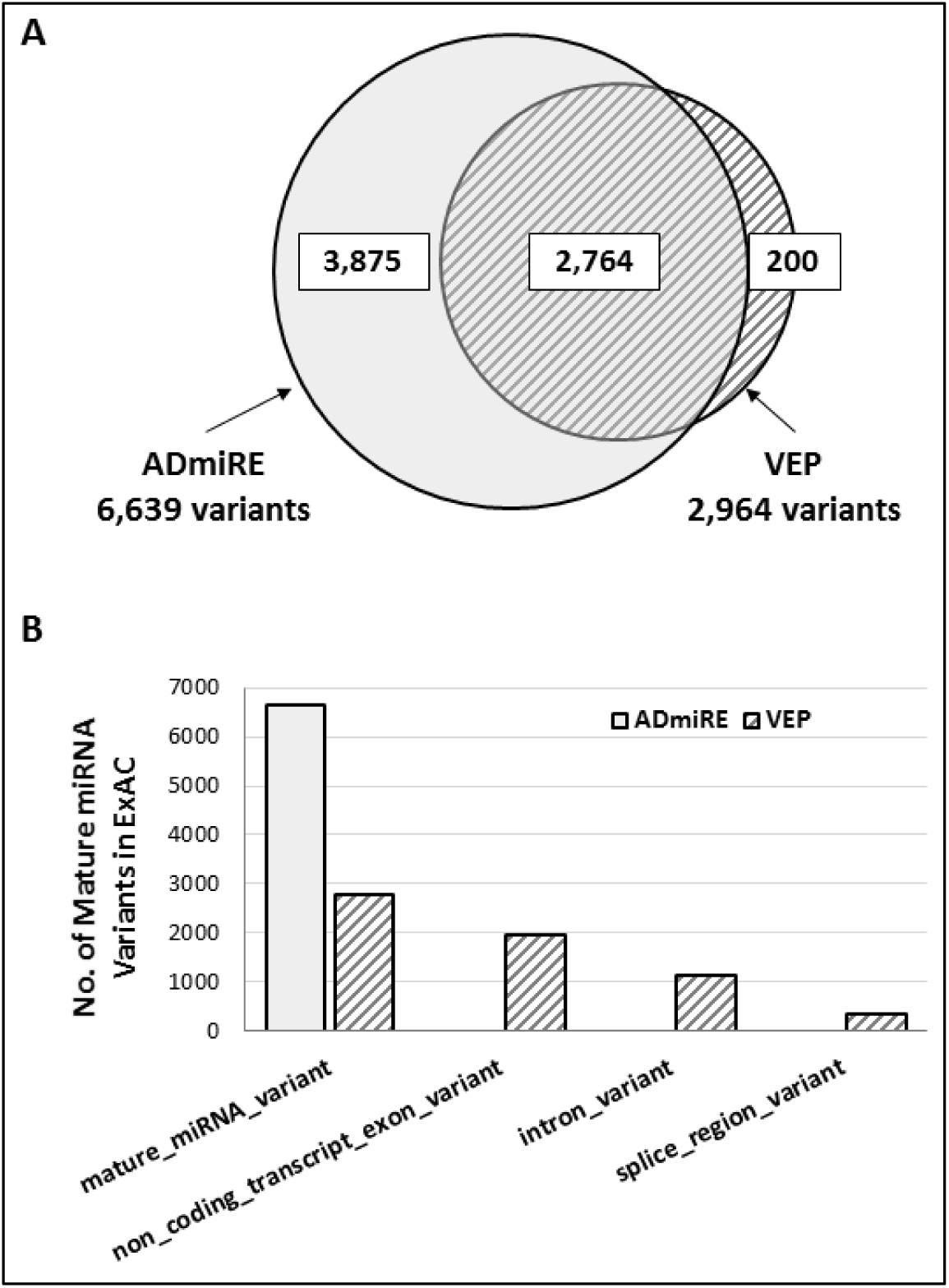
Evaluation of efficiency to detect mature miRNA variants by ADmiRE and VEP by re-annotation of ExAC variants. **A.** No. of mature miRNA variants annotated by ADmiRE (light gray) compared to VEP (hatched) annotations. **B.** VEP (hatched) ‘most damaging’ consequence categories for all mature miRNA variants detected by ADmiRE (light gray) from ExAC dataset.

### Pattern of variation in microRNAs in the gnomAD dataset

To develop a comprehensive landscape of miRNA variation, we applied ADmiRE workflow to the publicly available high-quality datasets of gnomAD-WGS and gnomAD-WES subsets for the re-annotation of variants within miRNA regions. The gnomAD-WGS dataset contains 11,757 precursor miRNA variants including the mature and seed domains and an additional 29,920 variants in the primary miRNA transcripts outside of the precursor hairpin regions (Table 1). We provide the complete list of annotated miRNA variants in the **Supplementary Data**.

**Table 1.** miRNA Variants identified using ADmiRE from 1000G and gnomAD Datasets.

| miRNA Domain | 1000G<br>n= 2,504 | gnomAD WGS<br>n= 15,496 | gnomAD WES<br>n= 123,125 |
| --- | --- | --- | --- |
| **Total** | **17,202** | **41,677** | **52,327** |
| **Seed** | 549 | 1,485 | 2,628 |
| **Mature** | 1,139 | 3,064 | 5,576 |
| **Precursor arm** | 3,029 | 7,208 | 11,575 |
| **Primary** | 12,485 | 29,920 | 32,548 |

We compared the minor allele frequency distribution of precursor miRNAs with the exonic, intronic, and intergenic regions and found miRNA variation resembles exonic variation and is significantly different than the intronic and intergenic variation (Mann-Whitney adjusted p-value <10^16^) (Figure 4A). We also observed that minor allele frequency distribution for miRNA variation is consistent with previously reported patterns for other types of variation including exonic variation with singletons representing over 50% of miRNA variants called by ADmiRE (Figure 4B) (Auton et al. 2015; Lek et al. 2016). We next assessed whether this resemblance of miRNA variation to the exonic regions is confounded by the genomic proximity of miRNAs to coding regions. We compared the miRNA variation between intragenic miRNAs located near or within the exonic regions (n=1,020, GENCODE v19) and the remaining intergenic miRNAs (n=858).

**Figure 4.**
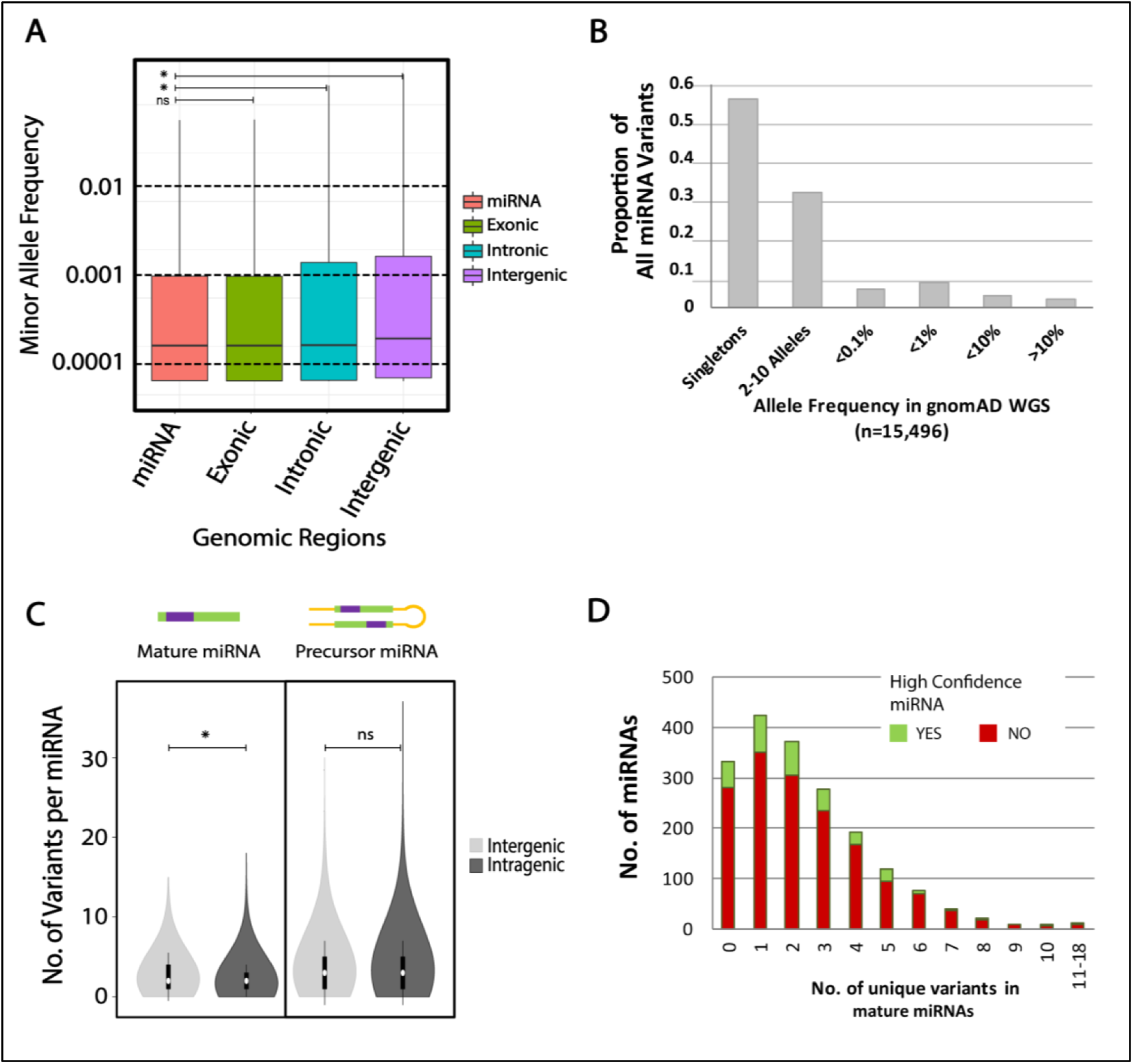
Patterns of miRNA variation in gnomAD-WGS (n=15,496) dataset. **A**. Distribution of minor allele frequency of all variants in log-scale across 4 different genomic regions, precursor miRNA (miRBase v21), exonic (GENCODEv19), intronic (GENCODEv19), and intergenic (remaining fraction of hg19 reference). **B.** The proportion of variants in precursor miRNA domains across the minor allele frequency spectrum, **C.** Variant distribution across mature and precursor miRNA domains compared between miRNAs in intragenic and intergenic regions (within 1kb of RefSeq genes). **D**. Number of miRNAs across each bin of number of unique mature domain variants. *-Mann-Whitney adjusted p-value <0.001, ns-not significant

Interestingly, we found that variant distribution of intergenic miRNAs was similar to that of intragenic fraction across precursor stem-loop domains however it was significantly different across the shorter mature domains likely driven by fewer mature domain variants in intergenic miRNAs (Mann-Whitney U test p-value <0.0001) (Figure 4C). We also find that the majority of mature miRNAs contain 1-2 variants in gnomAD, with a few outlier miRNAs accumulating very high sequence variation. Moreover, the ‘high confidence’ miRNAs are found to contain fewer variants compared to the remaining miRNAs (Figure 4D).

Sequences encoding domains essential for protein function are often intolerant to damaging sequence variation (Worth et al. 2009). There is a relative lack of such well-defined domains for the noncoding genome. However, for miRNAs, the seed and mature domains are the critical sequence domains for target gene recognition in miRNA-mediated regulation similar to critical protein domains. As expected, we observed that mature and seed domains contain significantly less variation than rest of the precursor domains and the flanking 100 bp primary domains in both the gnomAD-WGS and gnomAD-WES datasets (Figure 5A). Further assessment of the minor allele frequency distribution between high confidence and the remaining miRNAs demonstrates a significant difference in this distribution across all precursor domains with the distribution shifting towards rare variation in high confidence miRNAs (Mann-Whitney test p-value <0.001) (Figure 5B). We provide the first detailed landscape of miRNA variation across miRNA domains, position relative to gene regions and expression (as defined by a confidence score) from a large human population dataset. Overall, we observed that miRNA regions are under sequence constraint that is comparable to the protein domains in exonic regions independent of their surrounding genomic context, and the ‘high confidence’ miRNAs exhibit less variability compared to the remaining fraction.

**Figure 5.**
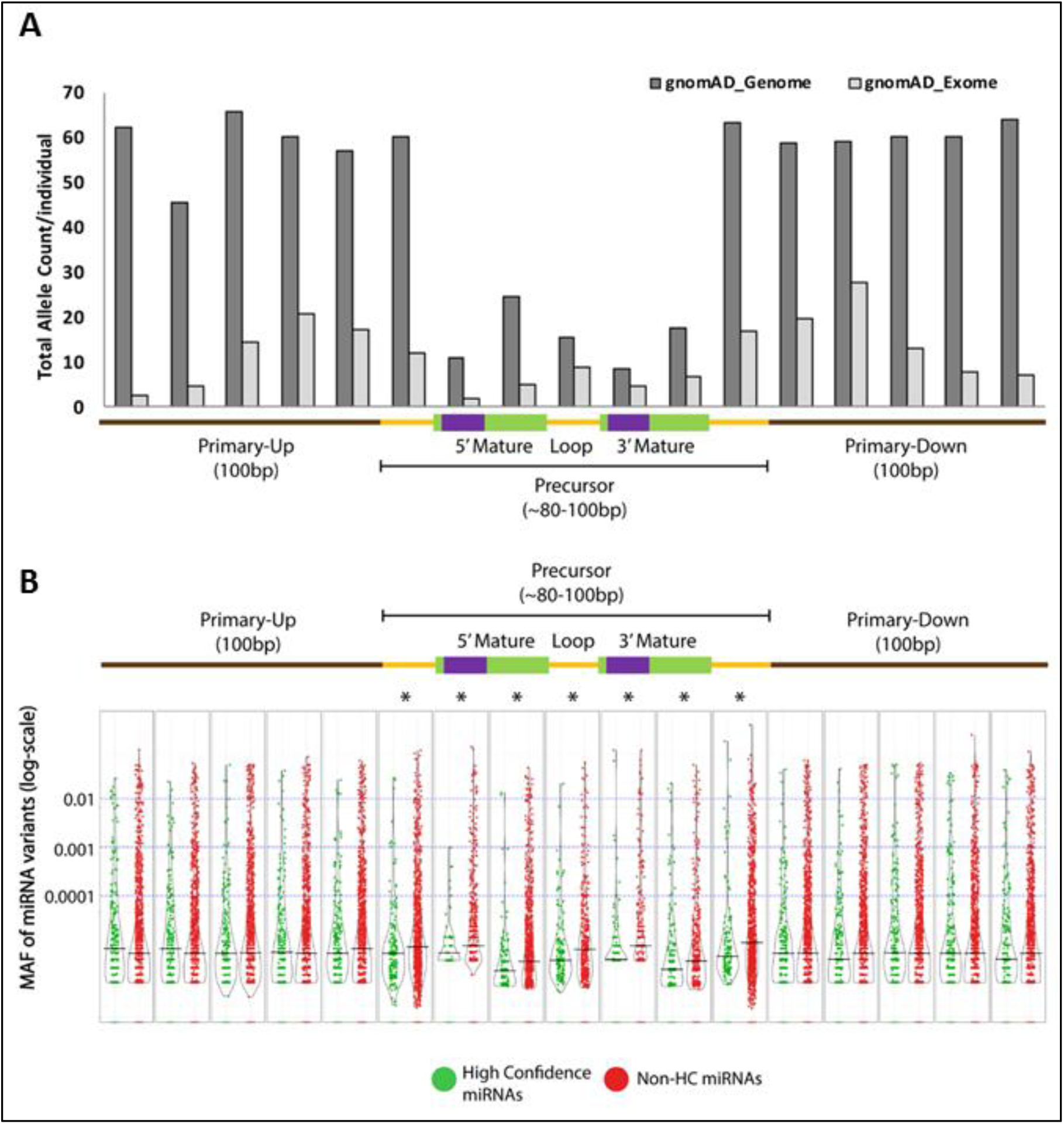
Distribution of miRNA variation in gnomAD across miRNA domains. **A.** Total number of miRNA variant alleles per individual re-annotated from the gnomAD-WGS (dark gray) and gnomAD-WES (light gray) subsets. The schematic shows each arm of primary miRNA transcript subdivided into five 20-bp regions (brown), precursor (yellow), mature (green) and seed (purple) domains, **B.** Violin plot of minor allele frequency distribution (log-scale) of miRNA variants from gnomAD-WGS subset normalized over domain length, median shown. Variants in ‘high confidence’ (green) and rest of the miRNAs (red) are plotted separately. *-Mann Whitney adjusted p-value <0.005

### Evaluation of miRNAs under constraint in gnomAD

Essential human genes are often intolerant to the damaging loss of function variation and exhibit constraint against such variation (MacArthur et al. 2012; Lek et al. 2015). We further examined which miRNAs are intolerant to sequence variation in the mature and seed miRNA domains in existing datasets. As most miRNAs have a short and uniform length of the mature domain (18-25 bp) we adopted a simple definition of a constraint. We first identified the 335 miRNAs that did not contain any mature domain variants in the gnomAD-WGS dataset (n=15,496) (Figure 4D). Further, we eliminated the miRNAs which contained a total of >10 alleles (approx. allele frequency >5*10^6^) in the gnomAD-WES dataset (n=123,136) resulting in 279 miRNAs highly constrained in humans (Supplementary Data). Gene ontology analysis of the experimentally identified targets for 279 constrained miRNAs showed significant enrichment for many essential cellular pathways such as regulation of apoptotic signaling, epithelial cell proliferation, cell motility and cellular migration (p-value < 10^-27^ -see Methods) (Supp. Table 2).

In contrast, we also evaluated the 45 miRNAs with greater than 4 seed domain variants depicting increased levels of variation, for example, miR-6087 with 18 mature domain variants and 12 seed domain variants. Some of these miRNAs may be artifacts of the miRNA detection tools so we assessed the putative biological impact of 15 seed domain SNVs from a curated list of 7 ‘high-confidence’ miRNAs (Supp. Table 3). We analyzed a common polymorphism from this list, miR-4707;rs2273626;G>U (MAF=40%) by gene ontology analysis using miRmut2Go webtool (Bhattacharya and Cui 2015). This algorithm predicts downstream targets for each of the alleles using TargetScan and miRanda tools, performs a gene ontology analysis for biological pathways and estimates the correlation of target pathways between the two alleles. This analysis suggested that this common polymorphism (rs2273626) was predicted to change >50% of the original targets of miR-4707 and generated a similar number of novel downstream target sites including many genes in the cellular signaling and cell communication pathways. Overall, we describe the frequency and distribution of sequence variation across all human miRNAs in the gnomAD dataset which allows us to identify highly constrained and highly variable miRNAs.

### Analysis of miRNA sequence conservation across 100 vertebrates

It is expected that a thorough understanding of the evolutionary conservation of miRNA sequences will further aid the evaluation of miRNA variation. We examined all human miRNAs using conservation scores generated by phyloP and phastCons scoring algorithms across 100 vertebrates. We divided the precursor miRNAs into groups, miRNAs with both 5’ and 3’ mature domains (n=931) and miRNAs with only one mature domain (n=946) and analyzed domain-wise sequence conservation of precursor miRNAs. We then performed hierarchical clustering of domain-wise conservation scores for each of these groups and identified 331 two-domain (Figure 6) and 103 single-domain (Supp. Fig. 2) conserved miRNAs based on top two clusters from the heatmap (Supplementary Data). Independently, we identified a similar proportion of conserved miRNAs (36% of 2,585) through the analysis of just the mature domains (Supp. Fig. 3).

**Figure 6.**
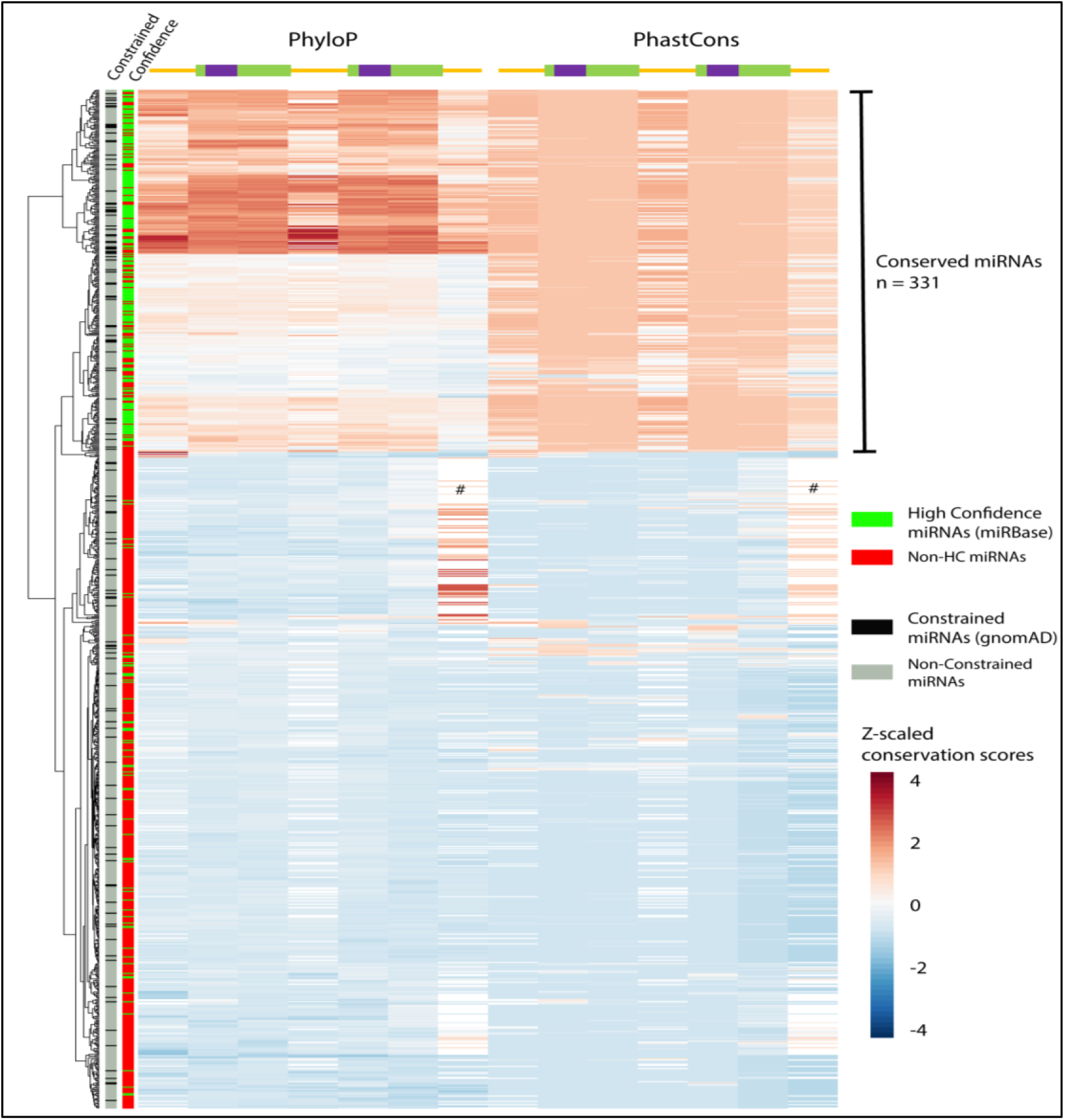
Heatmap of phyloP and phastCons 100way vertebrate conservation scores across precursor domains of miRNAs. phyloP and phastCons scores are centered and z-scaled across the precursor domains for each miRNA for which both the 5’ and 3’ mature domains are annotated by miRBase (931 miRNAs). On the left, hierarchical clustering of conservation across phyloP and phastCons scores for each miRNA. Each miRNA is annotated for whether it is a constrained miRNA (black) in human population dataset gnomAD, or not (grey). Each miRNA is also annotated whether it is a ‘high confidence’ miRNA (green) or otherwise (red). # -high homology in the 3’ precursor domain is the result of miRNAs that are immediately adjacent to coding exons.

As before, we confirm that the genomic context of intragenic and intergenic miRNAs does not confound the conservation of miRNAs as noted by non-significant difference in conservation scores for the two groups (Mann-Whitney U test, p-value = 0.23) (Supp. Fig. 4). However, we found a significant enrichment of the 279 constrained miRNAs in the conserved group of 434 miRNAs with an overlap of 65 miRNAs between the two groups (Chi-square p <0.0001).

High confidence’ miRNAs were also highly enriched in the group of highly conserved miRNAs (Chi-square p <0.0001). As described earlier, the ‘high confidence’ annotations were adopted to distinguish ‘real’ miRNAs from the non-miRNA reads detected by deep sequencing studies (Kozomara and Griffiths-Jones 2014). However, we also find that a few of the highly conserved miRNAs do not classify as ‘high confidence’ miRNAs. We investigated the expression of these non-high confidence yet highly conserved miRNAs across human tissues using miRmine database (Panwar et al. 2017). A few of the conserved miRNAs such as miR-21 and miR-124 indeed had a very low endogenous expression across all human tissues (Supp. Fig. 6A, 6B). A second group of miRNAs appear to be misclassified, for example, the non-high confidence miRNAs, let-7c and let-7i of the let-7 family exhibit high expression across all tissues, like that of the other ‘high confidence’ members of the let-7 family. (Supp. Fig. 6C, 6D). miRNAs such as miR-214 and miR-216 present an interesting case as these miRNAs are expressed at high levels in certain tissues while showing no expression in the remaining tissues. This tissue-specific expression can also be a possible cause of these miRNAs being misclassified (Supp. Fig. 6E, 6F). In conclusion, we have identified a group of highly conserved miRNAs across evolution that can be utilized for prioritizing the biological impact of miRNA variation. Further, we have shown a strong association of miRNAs with high expression in human tissues and conservation across vertebrates thus further providing a measure for predicting biological impact. Thus, three features of miRNA sequence (1) sequence constraint in human population datasets, (2) evolutionary conservation, and (3) ‘high confidence’ expression annotations are expected to further aid the interpretation of miRNA variation and expression patterns in disease datasets.

## Discussion

In this study, we have developed a dedicated miRNA variant annotation tool, ADmiRE, that both improves calling of miRNA sequence variants from VCF and BAM files and provides extensive miRNA and variant annotations developed here from the analysis of the large gnomAD population dataset and sequence conservation across evolution. ADmiRE can be accessed for calling miRNA variants from BAM files or re-annotation of existing set of variants in a number of ways (see Data Access). Using ADmiRE, we have identified approximately twice as many miRNA variants compared to a commonly used variant annotator, VEP, and show that VEP often prioritizes the protein-coding context (including introns) over miRNA context. We believe the use of ADmiRE workflow as an adjunct to existing annotation workflows will improve the detection of functional miRNA variation. To support this use, we have added the utility to use ADmiRE as a custom database to the VEP annotation tool. Not surprisingly, our analyses demonstrate that high coverage WGS is the ideal approach to comprehensively capture miRNA variation. However, given the large number of research and clinical WES efforts, we note that current exome capture platforms appear to capture approximately 50% of miRNAs in miRBase v.21 and these capture designs should be updated to improve evaluation of miRNA variation through WES analysis.

We evaluated the landscape of miRNA variation in 1,879 miRNAs (miRBase v21) from gnomAD, a publicly available collection of variation from high-quality 15,496 WGS and 123,136 WES samples as well as the smaller 1000G and UK10K projects. The pattern of sequence variation in miRNAs (SNP density and domain-level constraints) is very similar to that reported for protein coding exons. This was true for miRNAs found in intragenic regions and intergenic regions. In fact, intergenic miRNAs harbor fewer variants compared to the intragenic miRNAs and contain higher proportion of singleton variants. Thus, the properties of miRNAs described here are independent of proximity to protein-coding genes. We determined the per basepair variation across miRNAs domains and show that the seed and mature domains harbor much less variation compared to the rest of miRNA transcript. Domain level constraints are currently used by most existing tools for the prediction of biological impact of protein coding variants and the data provided here should improve these predictions for miRNA variation as prior efforts for non-coding RNAs have been challenging (Worth et al. 2009).

Identification of damaging variants in genes that are under constraint against loss of function variation facilitates the discovery of causal genetic loci for many rare Mendelian disorders (MacArthur et al. 2012). We have used the gnomAD datasets to generate a similar list of 279 constrained miRNAs. Variants in these constrained miRNAs are expected to have greater biological impact. We verified this in part by performing a gene ontology analysis of the experimentally identified targets of constrained miRNAs to find enrichment of cell proliferation, cell-cycle and cellular signaling pathways among others (Supp. Table 2). Conversely, there are a few common polymorphisms within seed sequences that are likely to substantially alter the pattern of target gene expression and may yield important population differences.

miRNA-mediated silencing pathways are conserved across eukaryotic lineages and many miRNA loci are also conserved across evolution (Altuvia et al. 2005; Chapman and Carrington 2007). However, there has not been a comprehensive evaluation of miRNA sequence conservation. In this study, we have determined that only 40% of human miRNAs (n=435 precursor, 920 mature) are highly conserved across 100 vertebrates. We further evaluated the overlap of constrained and conserved miRNAs with known disease mechanisms. We used KEGG pathway ‘MicroRNAs in Cancer’ to assess the proportion of miRNAs that are known to cause human diseases, such as cancer (Supp. Fig. 5). Interestingly, 145 of 150 miRNAs implicated across 10 types of cancer are highly conserved and 47 of these miRNAs are highly constrained for sequence variation (Table 2).

**Table 2.**
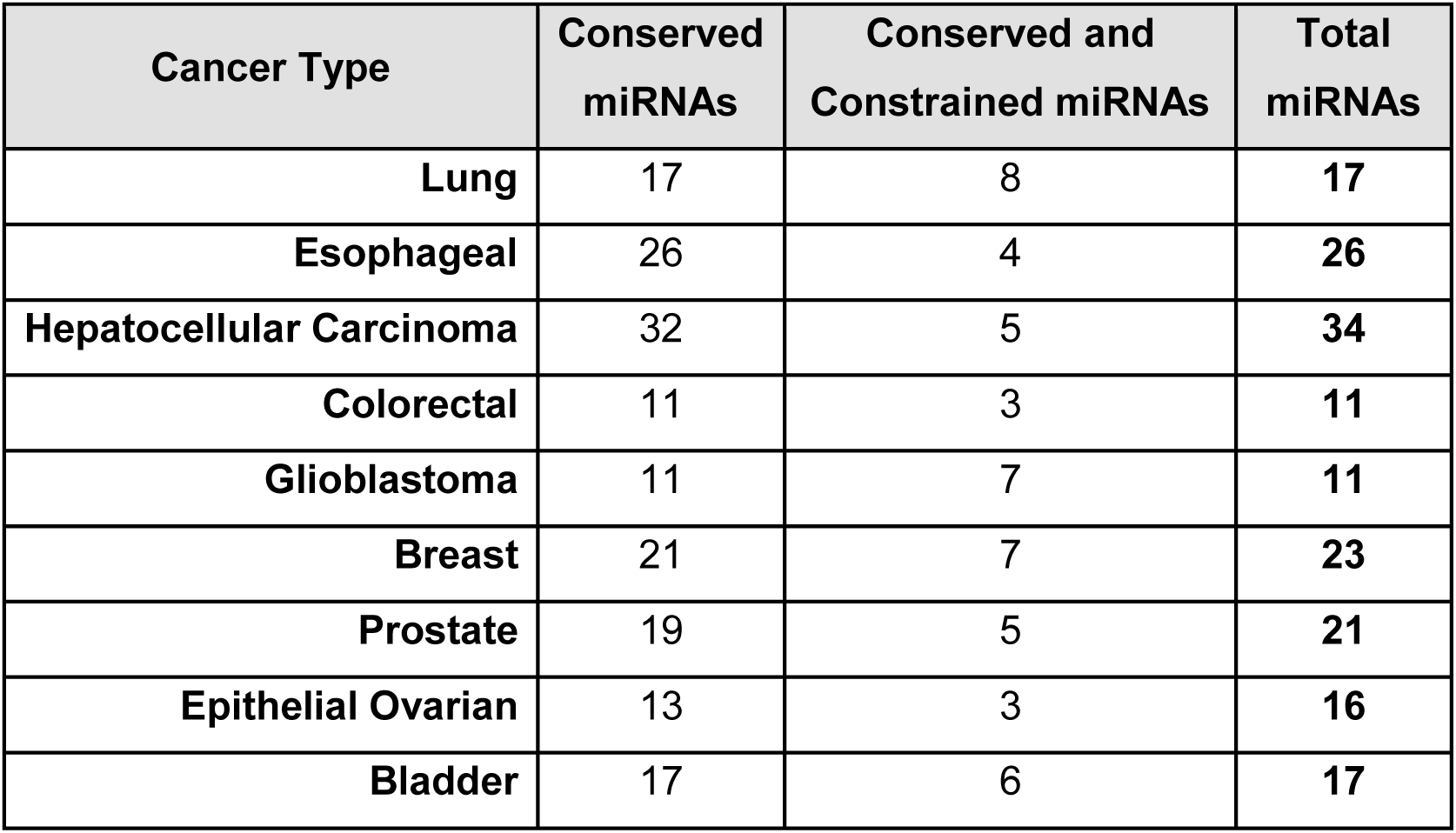
Number of conserved and constrained miRNAs in KEGG pathway, ‘miRNAs in Cancer’.

| **Cancer Type** | **Conserved miRNAs** | **Conserved and Constrained miRNAs** | **Total miRNAs** |
| --- | --- | --- | --- |
| **Lung** | 17 | 8 | **17** |
| **Esophageal** | 26 | 4 | **26** |
| **Hepatocellular Carcinoma** | 32 | 5 | **34** |
| **Colorectal** | 11 | 3 | **11** |
| **Glioblastoma** | 11 | 7 | **11** |
| **Breast** | 21 | 7 | **23** |
| **Prostate** | 19 | 5 | **21** |
| **Epithelial Ovarian** | 13 | 3 | **16** |
| **Bladder** | 17 | 6 | **17** |

Somewhat surprisingly, 60% of human miRNAs (including some that are constrained to variation) are not highly conserved across 100 vertebrates indicating their relatively younger evolutionary origin. Similar trend of evolutionary conservation is also observed in other multi-species comparisons across placental mammals and primates (Supp. Fig. 8). We noticed higher phyloP scores for primate-only alignments compared to those across 100 vertebrates, however, we did not observe many new conserved miRNAs within primates. Previously, researchers have shown that human-specific miRNAs are responsible for specialized human brain development (Arcila et al. 2014) and that coevolution of primate-specific miRNA-mRNA networks contributes to the increased primate cortex computational abilities (Londin et al. 2015).

In this study, we provide a detailed evaluation of the ‘high confidence’ annotations provided by miRBase (v21) catalogue of miRNAs. The purpose of these annotations is to distinguish bona fide miRNAs from putative misannotation of miRNA sequences by utilizing the deep sequencing RNA evidence supporting each miRNA. However, currently only 15% of miRNAs are classified as bona fide ‘high confidence’ miRNAs. We found that ‘high confidence’ miRNAs exhibit a higher sequence constraint in gnomAD dataset and are enriched in cluster of miRNAs conserved across 100 vertebrates. Interestingly, many constrained and conserved miRNAs lack the ‘high confidence’ annotations and many recent expression studies indicate their tissue-specific expression patterns. In this first detailed analysis of ‘high confidence’ miRNAs, we indicate the advantages and limitations of these annotations for the purpose of miRNA variant interpretation and suggest the need for ongoing updating of this group of miRNAs.

As reviewed in the Introduction, there have been several studies where single nucleotide variants in miRNAs have been identified in disease states. Interestingly, when we annotated these previously reported variants with the miRNA characteristics described in this study, we find that all 8 of the miRNAs harboring rare disease associated variants impact highly conserved, constrained and ‘high confidence’ miRNAs (Table 3). In this study, we have established a framework for improving the calling of miRNA variants from next generation sequencing datasets and enrich the annotations of these variants with experimentally validated targets of miRNAs and human disease associations from literature, in addition to the conservation, sequence constraint and confidence information developed using ADmiRE. This information will further guide the assessment of likely causal variation when exploring disease datasets such as those generated in the Mendelian Genomics and Undiagnosed Disease programs. Thus, ADmiRE provides the miRNA-focused annotations that can aid the identification and interpretation of biologically and medically relevant miRNAs when analyzing genome sequencing datasets.

**Table 3.** Sequence features for miRNAs previously described in human diseases.

| miRNA | Domain | Disease/<br>Cancer | Constrained<br>(gnomAD) | Conserved<br>(100 Vert.) | High<br>Confidence |
| --- | --- | --- | --- | --- | --- |
| miR-30c-1 <sup>1</sup> | Precursor | Breast | YES | YES | YES |
| miR-191 <sup>1</sup> | Precursor | Ovarian | YES | YES | YES |
| miR-128b <sup>2</sup> | Primary | ALL | YES | YES | YES |
| miR-16 <sup>3</sup> | Primary | CLL | YES | YES | YES |
| let-7e <sup>4</sup> | Primary | Prostate | YES | YES | YES |
| miR-17 <sup>1</sup> | Primary | Breast | YES | YES | YES |
| miR-96 <sup>5</sup> | Seed | Hearing Loss | NO * | YES | YES |
| miR-146a <sup>6</sup> | Mature | Cancer<br>Predisposition | NO * | YES | YES |

\*A single variant in gnomAD; <sup>1</sup>Kotani et al., 2010 <sup>2</sup>Calin et al., 2005 <sup>3</sup>Wu et al., 2008 <sup>4</sup>Shen et al., 2009 <sup>5</sup>Mencia et al., 2009 <sup>6</sup>Xu et al., 2011

## Methods

### Genomic Datasets

Single nucleotide variants (SNVs) and small insertions and deletions (InDels) from the genome aggregation database (gnomAD) consisting of 15,496 WGS samples and 123,136 WES samples were retrieved in the VCF format from the project’s download page (date accessed 2/28/2017). The gnomAD VCFs were reannotated using ADmiRE workflow with a runtime of <5 minutes per chromosome. For the development of the variant calling pipeline, WGS sequence alignments (BAM, n=2,504) and integrated variant callset (VCF) from the 1000 Genomes project dataset phase 3 release (1000G) were retrieved from the project’s FTP site (date accessed 1/3/2016) (Auton et al. 2015). The quality control metrics for the variant calling component of ADmiRE workflow were computed using of GATK variant evaluation tool, VariantEval. VariantEval uses an orthogonal genotyping platform (OMNI SNParray) to evaluate the accuracy of the variant caller. We observed a high genotype concordance between variants called by ADmiRE and those reported using the OMNI SNParray (>97%) and the expected Ti/Tv ratio of 2.26 (Supp. Table 1).For analysis of miRNA variation in additional publicly available sequencing dataset, the annotated variant files in the tab-separated format was retrieved from the Exome Aggregation Dataset (ExAC; date accessed 9/21/2016) and UK10K project (date accessed 2/21/2017) (The UK10K Consortium).

### MicroRNA Annotation Resources

We obtained the precursor stem-loop (1,881) and mature (2,815) miRNA coordinates (assembly GRCh38) for all human miRNAs from miRBase v21 and converted them to GRCh37 build using UCSC LiftOver tool (Kozomara and Griffiths-Jones 2014). Missing information between the two assemblies resulted in the final list of 1,878 precursor, and 2,785 mature miRNAs. We defined miRNA sequence domains as primary (100bp flanking the precursor), precursor, 5’ and 3’ mature, seed region, loop, and pre-5’ and pre-3’ flanking sequences (Figure 1A). We also retrieved a list of 295 ‘high confidence’ miRNAs from miRBase v21 defined by the following criteria (a) miRNAs with at least 10 reads mapping to each mature domain from RNA sequencing analyses, or (b) miRNAs with least 5 reads mapping to each mature domain and at least 100 reads mapping the entire precursor domain (Figure 1B) (Kozomara and Griffiths-Jones 2014). Additionally, we mined the literature and biological databases for information about associated diseases and experimentally validated target genes for all miRNAs and compiled data from 4 of the following databases: Human microRNA Disease Database (HMDD) (Li et al. 2014; Lu et al. 2008), PhnomiR (Ruepp et al. 2010), miRTarBase(Chou et al. 2016) and TarBase 7.0 (Vlachos et al. 2015). These annotations along with the miRNA sequence features described in this study for constrained and conserved miRNAs were compiled into a single database, <u>A</u>nnotative <u>D</u>atabase of <u>miR</u>NA <u>E</u>lements, ADmiRE (see section: **Data Access**).

### Cross-species miRNA conservation heatmap

We retrieved per base conservation scores for all miRNAs using UCSC table browser utility (date accessed 7/7/2016) for the following cross-species comparisons: Algorithms used were phyloP and phastCons across 100 vertebrates, 46 vertebrates, 46 placental mammals and 46 primates (Pollard et al. 2010; Siepel et al. 2005; Yang 1995; Felsenstein and Churchill 1996). For each conservation scoring system, phyloP and phastCons, scores were averaged across each miRNA domain resulting in 7 domain wise scores for phyloP and 7 for phastCons for each precursor miRNA. As the ranges of phyloP and phastCons scores differ widely (−13 to 10 for phyloP and 0-1 for phastCons) we normalized them across all miRNAs to a Z-score scale. The resulting matrix was hierarchically clustered using the default arguments of the aheatmap function as implemented in the NMF package in R (Kim et al. 2003).

### Statistical Analysis

All statistical analyses described in the study were performed using R statistical software (R-3.0.1). The variants in the exonic, intronic and intergenic regions did not follow a normal distribution. We used the Mann-Whitney test with a Bonferroni adjusted p-value (as implemented in the function pairwise.wilcox.test) for computing statistically significance of differences in the distribution of miRNA variants (Is this the variant distribution or frequency distribution) in the exonic, intronic, intergenic regions and intragenic regions (Figures 4A and 4B). To compare whether the distribution of variants and their minor allele frequencies differs among ‘high confidence’ and rest of the miRNAs, we used the same test for each miRNA domain (Figure 5B). To test whether ‘high confidence’ miRNAs and constrained miRNAs show enrichment in highly conserved clusters, the chi-square test was performed. To perform a gene ontology analysis of the constrained miRNA targets, we compiled a list of experimentally identified targets for each miRNA as per ADmiRE annotations and performed a gene ontology enrichment analysis using a webtool, Enrichr (Chen et al. 2013; Kuleshov et al. 2016).

## Data Access

ADmiRE workflow for annotation of user-supplied variant files: https://github.com/nroak/admire

## Acknowledgments

The work presented here was supported by Research grant RP10189 from the Cancer Prevention and Research Institute of Texas, NIH-R01-CA138836 and NHGRI 5 U01 HG007436-03 to SEP. The authors would like to thank the Genome Aggregation Database (gnomAD) team and the groups that provided exome and genome variant data towards this resource. A full list of contributing groups can be found at http://gnomad.broadinstitute.org/about. The authors would also like to thank Dr. Stephen Montgomery, Stanford University, for his comments on the manuscript.

## Supplementary Figures

**Supp. Fig. 1 miRNA region coverage across different sequencing platforms and annotations in WES datasets**

Analysis of 20 samples from 1000G dataset sequenced using all three platforms of WES, low coverage WGS and high coverage WGS at 5 different sequencing centers. Bedtools-coverageBed was used to compute the fraction of miRNA targets sequenced at corresponding read depth

**Supp. Fig. 2 Heatmap of phyloP and phastCons 100way vertebrate conservation for miRNAs with only 1 mature domain**

Hierarchical clustering of conservation across phyloP and phastCons scores for each of the 946 precursor miRNAs with only 1 mature domain.

**Supp. Fig. 3 Heatmap of phyloP and phastCons 100way vertebrate conservation across 2,785 mature miRNAs**

Hierarchical clustering of conservation across phyloP and phastCons scores for each of the 2,785 mature miRNAs. Each miRNA is annotated for constraint and a ‘high confidence’ annotations.

**Supp. Fig. 4 Comparing variant distribution between intragenic and intergenic conserved miRNAs.**

Number of variants in both mature and precursor domains from conserved miRNAs that resided within intragenic and intergenic regions. Mann Whitney U-test, ns-not significant

**Supp. Fig. 5 Sequence conservation and constraint of miRNAs altered in cancers** KEGG pathway “microRNAs in Cancer” colored according to the conserved and constraint annotations described in this paper

**Supp. Fig. 6 Analysis of highly conserved miRNAs that lack ‘high confidence’ annotations**

Using miRmine database to retrieve miRNA expression data for candidate miRNAs from top cluster of highly conserved miRNAs and the low confidence annotations.

**A.** Expression of a conserved, low confidence miRNA, miR-21, across all tissues. **B.** miR-124, a true low confidence miRNA with very little expression across all tissues. **C and D,** let-7 family with majority of ‘high confidence’ miRNA members except for let-7c and let-7i. **E and F.** miR-214 and miR-216 exhibit tissue specific expression.

**Supp. Fig. 7 Variant distribution across miRNA sequence domains**

Distribution of miRNA variants per individual identified using ADmiRE workflow. miRNA Variants were either recalled from 1000G BAMs (dark grey) or reannotated from 1000G VCF (light grey), ExAC variant table (dotted line) and UK10K variant table (solid line). The schematic below represents primary (brown), precursor (yellow), mature (green), and seed (purple) domains

**Supp. Fig. 8 Heatmap of conservation scores across precursor domains for miRNAs with both the mature domains expressed**

Conservation scores include phyloP and phastCons scores for 100way and 46way vertebrates, 46way mammals and 46way primate comparisons. Heatmap is ordered by hierarchical clustering across conservation scores for 100way vertebrate comparisons. On top is a schematic for miRNA domains such as precursor (yellow), mature (green), and seed (purple) domains. On the left, ‘high confidence’ and constrained miRNAs are annotated.

## Supplementary Tables

**Supp. Table 1 Summary of miRNA variants recalled from 1000G BAM and VCF files using ADmiRE Pipeline**

**A.** Distribution of miRNA variants analyzed using ADmiRE pipeline by

**i)** recalling variants from low coverage 1000G BAMs and

**ii)** re-annotating publicly released integrated variant callset.

**B.** QC metrics for jointly called variants from 1000G low coverage BAMs using ADmiRE pipeline.

**Supp. Table 2 Gene Ontology analysis for constrained miRNAs**

598 experimentally validated targets of 279 constrained miRNAs were analyzed for GO analysis using Enrichr webtool.

**Supp. Table 3 Seed domain SNVs in highly variable, ‘high confidence’ miRNAs**

List of SNPs from ‘high confidence’ miRNAs with greater than 4 seed domain SNPs.

## Supplementary Data

**Excel file: Supplementary_Data.xlsx**

a. Table of all human precursor miRNAs (miRBase v21, GRCh37) annotated with miRNA family names (miRBase v21), ‘high confidence’ (miRBase v21), constrained in gnomAD (this study), and conserved across 100 vertebrates (this study).
b. Complete list of miRNA variants in gnomAD-WGS dataset.
c. Complete list of miRNA variants in gnomAD-WES dataset.
d. Complete list of miRNA variants in 1000 Genomes (phase 3) dataset.
e. Complete list of miRNA variants in ExAC dataset.
f. Complete list of miRNA variants in UK10K dataset.

